# P4HA1 sustains endocrine resistance in luminal breast cancer via regulation of cellular stress

**DOI:** 10.64898/2026.09.30.755755

**Authors:** Dalton Hironaka, Baorui Kang, Dongsung Kim, Haichang Li, Rachel Schiff, Ren Xu, Gaofeng Xiong

**Affiliations:** Department of Veterinary Biosciences, College of Veterinary Medicine, The Ohio State University, Columbus, OH 43210, USA; Molecular, Cellular and Development Biology Program, The Ohio State University, Columbus, OH 43210, USA; OSU Nutrition Graduate Program, The Ohio State University, Columbus, OH 43210, USA; James Comprehensive Cancer Center, The Ohio State University, Columbus, OH 43210, USA; Baylor College of Medicine, Houston, TX 77030, USA; Department of Radiation Oncology, University of Arkansas for Medical School, Little Rock, AR 72205, USA; Department of Pharmacology and Nutritional Sciences, College of Medicine, University of Kentucky, Lexington, KY 40536, USA

**Author notes:** Corresponding author: Gaofeng Xiong, 1900 Coffey Rd, VMAB 460. Columbus, OH 43210. These authors contributed equally to this work (Dalton Hironaka, Baorui Kang).

**Keywords:** Prolyl 4-hydroxylase, endocrine-resistant, ER stress, ROS

## Abstract

The hormone receptor-positive luminal subtype accounts for approximately 70% of all diagnosed breast cancer cases, with endocrine regimens serving as the first-line therapy. However, 30% to 50% of luminal breast cancer patients acquire therapeutic resistance with five to ten years of treatment. Understanding the mechanisms of endocrine resistance and identifying new therapeutic targets remain urgent priorities. Here, we demonstrate that expression of the alpha subunit of prolyl 4-hydroxylase (P4HA1), a key collagen-modifying enzyme, is significantly upregulated in endocrine therapy-resistant luminal breast cancer cells compared to parental endocrine therapy-sensitive cells. Clinically, elevated P4HA1 expression correlates with shortened relapse-free survival in patients with luminal breast cancer. Genetically knocking down P4HA1 or pharmacologically inhibiting its activity successfully restores the sensitivity of endocrine-resistant luminal breast cancer cells to hormone therapy. Mechanistically, we reveal that combining P4HA1 inactivation with endocrine treatment dramatically enhances endoplasmic reticulum (ER) stress-related gene expression. Consistently, treating resistant cells with an exogenous ER stress inducer similarly enhances sensitivity to hormone therapy. Furthermore, we demonstrate that ER stress induced by P4HA1 inactivation is molecularly mediated by an accumulation of reactive oxygen species (ROS). Together, these findings uncover a novel, non-canonical role for P4HA1 as a critical mediator linking endocrine resistance to cellular stress adaptations, positioning P4HA1 as a promising therapeutic vulnerability in endocrine-resistant breast cancer. P4HA1 serves as a necessary role in maintenance resistant state, making it a viable target for combination therapies to overcome endocrine resistance.

## Background

Collagen prolyl 4-hydroxylase (P4H), a member of the Fe^2+^ and alpha-ketoglutarate (α-KG)- dependent dioxygenase family, stoichiometrically utilizes molecular oxygen and α-KG to hydroxylate proline residues, releasing CO_2_ and succinate as byproducts [1]. The primary function of C-P4H is to hydroxylate proline residues at carbon 4 (C-4) within the Y-position of the -X-Y- Gly- triplet sequence motif in procollagen peptides. This key post-translational modification is critical for modulating protein folding and structural stability in the endoplasmic reticulum (ER) [1] [2]. Three P4H isoforms (P4H1, P4H2 and P4H3) have been identified in mammalian cells [2]. P4H1 is the major isoform that contributes to the majority of the P4H activity [3]. Structurally, P4H is a tetrameric enzyme composed of two α subunits (P4HA) and the two β subunits (P4HB). The α subunit is translated on ribosomes along the rough ER and subsequently binds to the β subunit, which is constitutively present in the ER lumen at a much higher level [4]. P4HA1, the alpha subunit of P4H1, is responsible for both peptide binding and catalytic activity [2]. Notably, P4HA1 is overexpressed in the majority of solid cancers ranging from breast cancer and colon cancer to melanoma and esophageal cancer [5, 6].

Luminal A, Luminal B, HER2-positive, and triple-negative breast cancer (TNBC) represent four major molecular subtypes of breast cancer. While previous studies have demonstrated that P4HA1 expression is significantly induced in TNBC, and its high expression is associated with poor survival in TNBC patients, less is known about its role in other subtypes. Cancer cell-derived P4HA1 plays an important role in cancer progression and metastasis, which is primarily mediated by hydroxylation during collagen synthesis [7, 8]. The hormone receptor-positive luminal subtype breast cancer accounts for about 70% of all breast cancer diagnoses. For these patients, endocrine therapies including selective estrogen receptor modulators (SERMs, such as tamoxifen), selective estrogen receptor degraders (SERDs, like fulvestrant) and specific aromatase inhibitors, are the first-line therapies [9]. Although endocrine therapies have significantly reduced mortality rates over recent decades [10], approximately 30% to 50% of luminal breast cancer patients eventually develop acquired resistance to endocrine therapy after five to ten years of treatment. Consequently, therapeutic resistance and lethal recurrence remain the primary drivers of mortality in breast cancer patients [11–13].

Acquired or *de novo* resistance to endocrine therapy is a consequence of complicated molecular alterations that allow tumor growth irrespective of estrogen presence. Several chemicals are being investigated to overcome resistance by interfering with signaling pathways activated in luminal breast cancer cells, such as the growth factor receptor/PI3K/AKT/mTOR signaling pathway, or the CDK4/6 signaling pathway [14]. Some are currently being tested in clinical trials, such as AKT inhibitors, ER partial agonists, next generation SERDs, EGFR/HER2 dual inhibitors, and HDAC inhibitors [15]. Currently, FDA-approved therapeutic strategies to improve the outcome of endocrine resistance in patients include inhibitors against PI3K, mTOR, ER proteolysis target chimeras (PROTACs) and CDK4/6 inhibitors [16, 17]. The treatment landscape for advanced luminal breast cancer patients has shifted toward combination strategies incorporating endocrine therapy with these targeted inhibitors, such as CDK4/6 inhibitors, which have become the standard of care. However, a substantial proportion of patients still fail to respond to current combination therapies, and resistance to these regimens remains a major clinical challenge. Therefore, a critical need persists in developing more effective therapeutic strategies capable of overcoming resistant contexts.

In this study, we demonstrate that increased expression of P4HA1 is associated with resistance to endocrine therapies, including tamoxifen and fulvestrant, in estrogen receptor positive luminal breast cancer cells. Inactivation of P4HA1 restored the sensitivity of resistant cells to hormone therapy. Targeting P4HA1 in endocrine-resistant cells was accompanied by enhanced ER stress-related gene expression and markedly elevated reactive oxygen species (ROS) levels. These findings reveal a previously unrecognized role of P4HA1 in contributing endocrine resistance through regulation of cellular stress networks and identify P4HA1 as a potential therapeutic target crucial to reducing recurrence in luminal breast cancer patients.

## Methods

### Cell lines, cell culture, and treatment

The human breast cancer cell line MCF7 (Cat# HTB-22, RRID:CVCL_0031), T47D (Cat# HTB-133, RRID:CVCL_0553) and human embryonic kidney cell line HEK293FT were obtained from American Type Culture Collection, tamoxifen-resistant T47D/TR-1 cell line was purchased from CancerTools.org (Cat # 152108, RRID:CVCL_1D36). The human breast cancer cell line LCC9, a variant of the MCF7 cell line, was a generous gift from Dr. Robert Clarke at Georgetown University. It was initially selected through *in vivo* passage in ovariectomized athymic nude mice, followed by *in vitro* selection with fulvestrant (from 10 pM to 1 μM). It represents a stable fulvestrant-resistant cell model that also exhibits complete cross-resistance to tamoxifen, despite no prior exposure to tamoxifen [18]. The MCF7/EndR cell line was established by continuous exposure of tamoxifen-resistant cells MCF7/TamR to fulvestrant (1 μM) for four months [19]. MCF7/TamR cells (from Dr. Rachel Schiff, Baylor College of Medicine) were previously generated through estrogen deprivation and continuous tamoxifen treatment for 6 months [20]. MCF7 and T47D are estrogen receptor-positive breast cancer cell lines that are highly sensitive to endocrine therapies and serve as endocrine-sensitive parental models, whereas MCF7 variants LCC9 and MCF7/EndR, and T47D variant T47D/TR-1 cell lines all serve as endocrine-resistant models.

MCF7, T47D, and MCF7/EndR cells were maintained in Dulbecco’s Modified Eagle’s Medium (DMEM) (Sigma Aldrich, D6429) with 10% fetal bovine serum (FBS) (Sigma Aldrich, F2442), 100 units penicillin and 0.1 mg streptomycin/mL (Sigma, P4333). HEK293 FT cells were maintained in DMEM (Sigma Aldrich, D6429) with 10% FBS (Sigma Aldrich, F2442), 1% MEM Non-essential Amino Acid Solution (Sigma Aldrich, M7145), 1 mM Sodium pyruvate solution (Sigma Aldrich, S8636), 2 mM L-Glutamine (Sigma Aldrich, G7513), 100 units penicillin and 0.1 mg streptomycin/mL (Sigma, P4333). LCC9 cells were maintained in Improved MEM (IMEM) (Corning, 10-026-CV) with 10% FBS, 100 units penicillin and 0.1 mg streptomycin/mL. T47D/TR-1 cells were maintained in RPMI 1640 (Sigma, R8758) with 10% FBS, 2 mM L-Glutamine, 8 μg/mL Insulin (Sigma Aldrich, I6634). All the cells were cultured in a humidified incubator at 37°C with 5% CO_2_ and 95% O_2_. All cells were treated with Plasmocin™ (Invivo Gen, ant-mpt-1) to eliminate and prevent mycoplasma contamination. Reagents used in luminal breast cancer cells treatments: 4-hydroxytamoxifen (Sigma, H6278); fulvestrant (Sigma, Y0001399); ethyl-3, 4-dihydroxybenzoic acid (DHB) (Sigma, 37580-25G-F); 1, 4-dihydrophenonthrolin-4- one-3-Carboxylic acid (1,4-DPCA) (Cayman, 331830-20-7); tunicamycin (Sigma, T7765); *N*- Acetyl-L-cysteine (NAC) (Sigma, BP907).

### Lentivirus-mediated knockdown or ectopic gene expression

Human *P4HA1* knockdown plasmids sh*P4HA1* (Clone ID: TRCN0000303933, target sequence: CCTGTGGTGTCTCGAATTAAT; Clone ID: TRCN0000303934, target sequence: ATTTGAGCTATGCGGTATATC) and human DDIT3 knockdown plasmid sh*DDIT3* (Clone ID: TRCN0000364328, target sequence: CTGCACCAAGCATGAACAATT) were purchased from Sigma. pCDH1-P4HΑl-Flag [8] was used to introduce ectopic P4HA1 expression. HEK293 FT cells were transfected with control vector, or gene knockdown or overexpression plasmid, together with packaging lentivectors using FuGENE® HD Transfection Reagent (Promega, E2312). Culture supernatants containing viral particles were collected 48 h after transfection. Human breast cancer cells were infected with lentivirus and stable-silenced cell lines were selected by puromycin for at least one week.

### Immunoblotting analysis

Cultured cells were lysed in 2% SDS in PBS buffer containing phosphatase and protease inhibitor cocktails (EMD Millipore, 539131). Protein concentration was measured using Pierce™ BCA Protein Assay Kit (Thermo Fisher Scientific, 23227). Equal amounts of protein lysates were subjected to SDS gel electrophoresis, electroblotted onto PVDF membranes (Bio-Rad, 1620177), and immunoblotted with primary antibodies (anti-P4HA1, Proteintech, 12658-1-AP, RRID:AB_2283162; anti-BIP Proteintech, 11587-1-AP, RRID:AB_2119855; anti-CHOP, Proteintech, 15204-1-AP, RRID:AB_2292610; anti-IRE1, Proteintech, 27528-1-AP, RRID:AB_2880899; anti-ATF4, Proteintech, 10835-1-AP, RRID:AB_2058600; anti-PERK, Proteintech, 24390-1-AP, RRID:AB_2879521; anti-Tubulin hFAB Rhodamine, Bio-Rad, 12004165, RRID:AB_2884950) at 4°C overnight, and StarBright Blue 700 Goat Anti-Rabbit IgG (Bio-Rad, 12004162, RRID:3712168) or StarBright™ Blue 520 Goat Anti-Mouse IgG (Bio-Rad, 12005867, RRID:AB_2934034) secondary antibodies for 2 hours at room temperature. Membranes were exposed using (Bio-Rad, ChemiDoc MP, RRID:SCR_019037). Western blotting results were quantified by using Image J analysis software.

### RNA-Seq Microarray analysis

Total cellular RNA samples of shRNA control and *P4HA1*-silenced LCC9 cells treated with or without tamoxifen were extracted using the RNeasy mini kit (Qiagen, 74106). RNA was quantified by measuring optical density at 260 nm, and quality was verified by agarose gel electrophoresis. NEBNext Ultra Directional RNA Library Prep Kit (New England BioLabs, Ipswich, MA) was used for library preparation. To analyze differential gene expressions, sequence reads were aligned to the genome by using a standard Illumina sequence analysis pipeline, which was analyzed by The Laboratory for Statistical Genomics and Systems Biology at the University of Cincinnati. DAVID analysis was performed to identify the differentially the expressed signaling pathways.

### Quantitative Reverse transcription-polymerase chain reaction (RT-qPCR)

Total RNA was extracted from shRNA control and *P4HA1*-silenced LCC9 cells treated with or without tamoxifen using TRIzol reagent (Invitrogen, 15-596-018). cDNA was synthesized using the SuperScript™ IV First-Strand Synthesis System (Invitrogen, 18091050) from 2.0 μg RNA samples. Synthesis of cDNA was performed according to the manufacturer’s instructions. RT-qPCRs were carried out with SYBR green PCR master mix reagents using a QuantStudio™ 3 Real-Time PCR System (Thermo Fisher Scientific, A28567, RRID:SCR_018712). Thermal cycling was conducted at 95°C for 30 sec, followed by 40 cycles of amplification at 95°C for 5 sec, 55°C for 30 sec, and 72°C for 15 sec. The relative quantification of gene expression for each sample was analyzed by the △ threshold cycle (△△CT) method. Information of primers used for amplification of ER stress pathway genes and 18S rRNA is in Table 1.

**Table 1:**
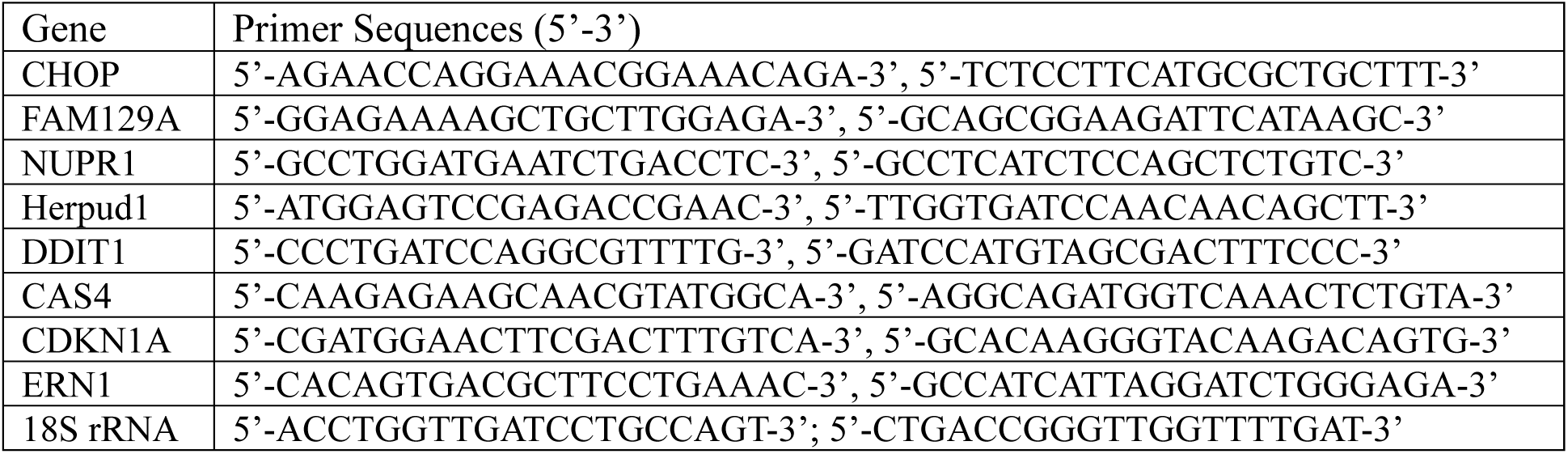
Primers used in RT-qPCR.

### CCK8 Cell viability analysis

Luminal breast cancer cells were seeded to a 96- well cell culture plate (3000 cells per well). After 24 h, cells were treated with P4H inhibitors (DHB or 1,4-DPCA), tamoxifen, fulvestrant, or a combination of a P4H inhibitor with tamoxifen/fulvestrant by adding each to the cell culture medium. After a 2-day treatment, 10 μL of the CCK-8 solution was added to each well of the plate, according to the manufacturer’s instruction for the Cell Counting Kit-8 (Sigma-Aldrich, 96992). The plate was incubated for 1 - 3 h at 37 °C in the cell incubator, then read on a Spectra MRTM microplate spectrophotometer (Dynex Technologies) at OD450 nm. For each cell clone, the OD450 values in the drug treatment group were normalized to the value in the control group.

### Cell proliferation analysis

Cell proliferation was examined using the Click-iT™ EdU Cell Proliferation Kit for Imaging, Alexa Fluor™ 488 dye (Invitrogen, C10337). According to the manual, EdU solution was added to the cell cultured medium at a final concentration of 10 μM and incubated for 8 h at 37 °C. After incubation, medium was removed from the cells and fixed with 4% PFA at room temperature for 15 min, followed by 0.5% Triton® X-100 permeabilization. Click-iT® reaction cocktail was incubated with the cells at room temperature for 30 min to detect EdU-positive cells. DAPI solution was incubated for 2 min, and pictures were captured using a Nikon Eclipse Ts2 microscope. EdU positive ratio was quantified by counting EdU and DAPI positive cells.

### Cell apoptosis analysis

Cell apoptosis assay was performed using Dead Cell Apoptosis Kits with Annexin V for Flow Cytometry (Invitrogen, V13242). According to the manual, 3–4 days after treatment with tamoxifen or fulvestrant in combination with P4H inhibitors, LCC9 or MCF7/EndR cells were harvested and incubated with FITC Annexin V and 1 μg/mL propidium iodide (PI) at room temperature for 15 min, and analyzed by flow cytometry with Becton Dickinson LSR II or Attune NxT flow cytometer (Thermo Scientific). Data were analyzed by FlowJo software.

### ROS level analysis

According to the manual of CellROX Deep Red Reagent (Invitrogen, C10422), the CellROX® Deep Red reagent was added to the cell cultured medium at a final concentration of 5 μM and incubated for 30 min at 37 °C. After 30 min incubation, culture medium was removed, and cells were washed with PBS three times. Cells were fixed with 4% PFA for 15 min. and images were taken with a Nikon Eclipse Ts2 microscope. Fluorescence intensity was quantified using the NIS Elements AR software.

### Immunohistochemical staining

Xenograft tumor sections were deparaffinized and rehydrated through submerging successively in 100% alcohol, 95% alcohol, 70% alcohol and then PBS solutions. Endogenous peroxidase was blocked by incubation with 3% H_2_O_2_ for 20 min. At the antigen retrieval step, slides were steamed in citrate sodium buffer for 30 min. Slides were blocked with Avidin/Biotin Blocking Kit (Vector Laboratories, SP-2001) and 5% goat serum, incubated with primary antibodies (anti-P4HA1, Proteintech, 12658-1-AP, RRID:AB_2283162; anti-Ki67, Cell Signaling Technology #9027, RRID:AB_2636984) at 4°C overnight, and then the sections were incubated with Biotinylated Goat Anti-Rabbit IgG Antibody (Vector Laboratories, BA-1000, RRID:AB_2313606) at room temperature for 60 min. After incubation with Streptavidin, Horseradish Peroxidase (Vector Laboratories, SA-5704, RRID:AB_2336692) at room temperature for 30 min, slides were stained with substrate diaminobenzidine (DAB, Vector Laboratories, SK-4100) and counterstained with hematoxylin. Images were captured using Olympus BX43F Microscope, and Ki67 positive ratio was quantified using Image J. Protein levels of P4HA1 expression in human breast cancer tissues were determined using tissue microarray (TMA) (BC081116f and BR1141b, TissueArray.com). Each case in the TMA is represented by three separate tissue cores. IHC scoring results from each of these three tissue cores were averaged to produce a final score for each case. P4HA1 immunohistochemical staining was scored according to a standard semi-quantitative scale as follows: negative (0), weak (1), moderate (2), and strong (3).

### *In vivo* Xenograft experiments

Eight-week-old female SCID mice (Taconic, ICRSC-F, RRID:IMSR_TAC:ICRSC) were subcutaneously implanted with 17β-estradiol pellet (0.72 mg, 60-day release, Innovative Research of American) under isoflurane anesthesia. One day after 17β-estradiol pellet implantation, parental LCC9 cells, shcontrol, or sh*P4HA1* LCC9 cells were harvested and injected into the 4^th^ mammary fat pad at 1 x 10^6^ cells/point. Tumors were measured with a caliper twice per week to analyze tumor growth. When tumors reached 60∼70 mm^3^, mice implanted with parental LCC9 cells were randomly grouped and treated with vehicle control or 40 mg/kg DHB intraperitoneal injection daily or 250 mg/kg fulvestrant subcutaneous injection weekly; mice implanted with shcontrol or sh*P4HA1* LCC9 cells were randomly grouped and treated with vehicle control or 250 mg/kg fulvestrant subcutaneous injection weekly. At the experimental endpoint, tumors were harvested and fixed with a 10% formalin solution for the paraffin-embedded section for immunohistochemical staining. All the procedures during our study were conducted within the guidelines of the Division of Laboratory Animal Resources at the University of Kentucky (Protocol # 2019-3371).

### Kaplan-Meier survival analysis

To address the clinical relevance of enhanced P4HA1 expression, we assessed the association between P4HA1 mRNA levels and patient survival using the published dataset generated from 1491 human luminal B breast cancer tissue samples. Tumor samples were equally grouped into low and high P4HA1 expression based on the mRNA levels. Significant differences in overall survival time were assessed with the Kaplan–Meier survival analysis [21].

### Statistical analysis

Results were reported as mean ± standard error of the mean (SEM), and the significance of difference was assessed by independent Student’s *t*-test or one-way analysis of variance (ANOVA) with GraphPad Prism software (RRID:SCR_002798). P < 0.05 is considered statistical significance, and P < 0.01 represented sufficient statistical significance.

## Results

### P4HA1 is associated with endocrine resistance in luminal breast cancer

Using human breast cancer samples from tissue microarrays (TMA), we observed that P4HA1 expression was markedly upregulated in breast cancer tissue compared to adjacent normal mammary gland tissue (Fig. 1*A,* 1*B*, Fig. S1*A*). High expression of P4HA1 was associated with poor survival in lymph node-positive and grade 3 patients, but not in lymph node-negative or lower grades patients (Fig. 1*C*, Fig. S1*B*). These clinical data indicate that elevated P4HA1 expression correlates with breast cancer malignancy. Interestingly, we found that strong P4HA1-positive staining was significantly enriched in Luminal B compared with Luminal A breast cancer (Fig. 1*A*, 1*B*). Kaplan-Meier survival analysis revealed that increased P4HA1 was associated with shorter relapse-free survival in patients with Luminal B breast cancer, but not in those with Luminal A (Fig. 1*C*). This highlights the clinical significance of P4HA1 in Luminal B subtype. While both subtypes are hormone receptor-positive breast cancer, Luminal B tumors are characterized by higher cell proliferation rate (e.g., elevated Ki67) and tend to respond less effectively to hormone therapy, leading to a higher risk of recurrence and poorer overall prognosis [22]. Our prior report reveals that P4HA1 promotes chemoresistance in TNBC [8], therefore, we hypothesized that high expression of P4HA1 may similarly contribute to endocrine resistance and recurrence in luminal breast cancer.

**Figure 1.**
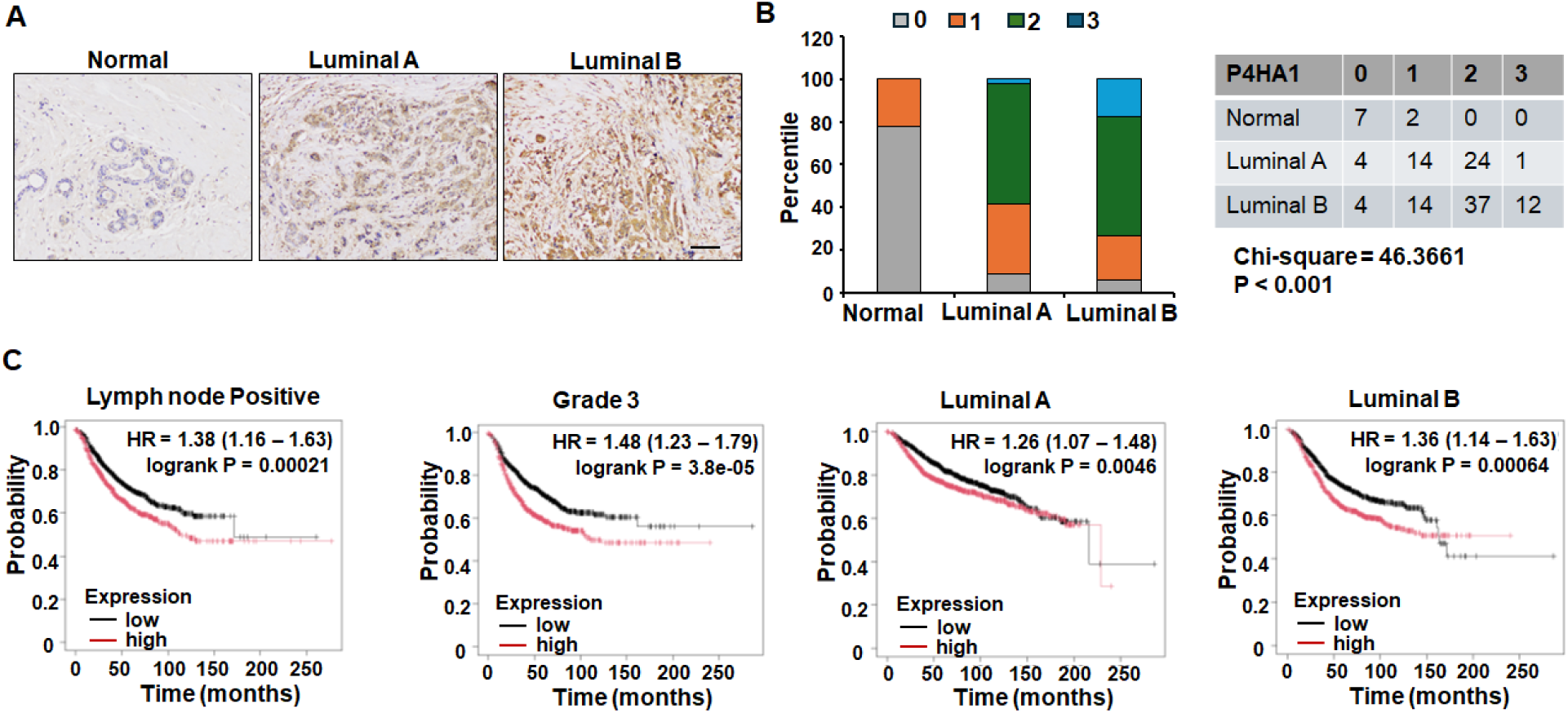
P4HA1 expression is associated with malignancy in luminal breast cancer patients. (A) P4HA1 IHC staining pictures and (B) staining intensity quantification in human mammary normal tissue (n=9), human Luminal A (n=43) and Luminal B (n=67). P4HA1 immunohistochemical staining was scored as follows: negative (0), weak (1), moderate (2), strong (3). Scale bar: 50 μm. (C) Kaplan Meier analysis of survival in lymph node positive (n=1656), grade 3 (n=1300), human Luminal A (n=43) and Luminal B (n=67) breast cancer patients with different levels of P4HA1 protein levels.

Next, we assessed P4HA1 expression across the endocrine-sensitive parental cell lines (MCF7 and T47D) and their endocrine-resistant derivatives (LCC9, MCF7/EndR, and T47D/TR-1) [18, 19]. Compared with parental MCF7 cells, MCF7 variants LCC9 and MCF7/EndR cells both exhibited resistance to the standard endocrine agents for luminal breast cancer patients: tamoxifen and fulvestrant (Fig. 2*A*, 2*C*). Compared with parental T47D cells, T47D/TR-1 cells displayed resistance to tamoxifen (Fig. 2*B,* 2*D*, Fig. S2*A*). Quantitative analysis revealed that P4HA1 protein levels were significantly elevated in all three endocrine-resistant luminal breast cancer cells (LCC9, MCF7/EndR, and T47D/TR-1) compared to their respective endocrine-sensitive parental counterparts (Fig. 2*E*, 2*F*, Fig. S2*B*). This upregulation was further confirmed at the mRNA level (Fig. 2*G*). Collectively, these luminal breast cancer cell line models and clinical findings link P4HA1 expression to endocrine resistance and recurrence in luminal B breast cancer.

**Figure 2.**
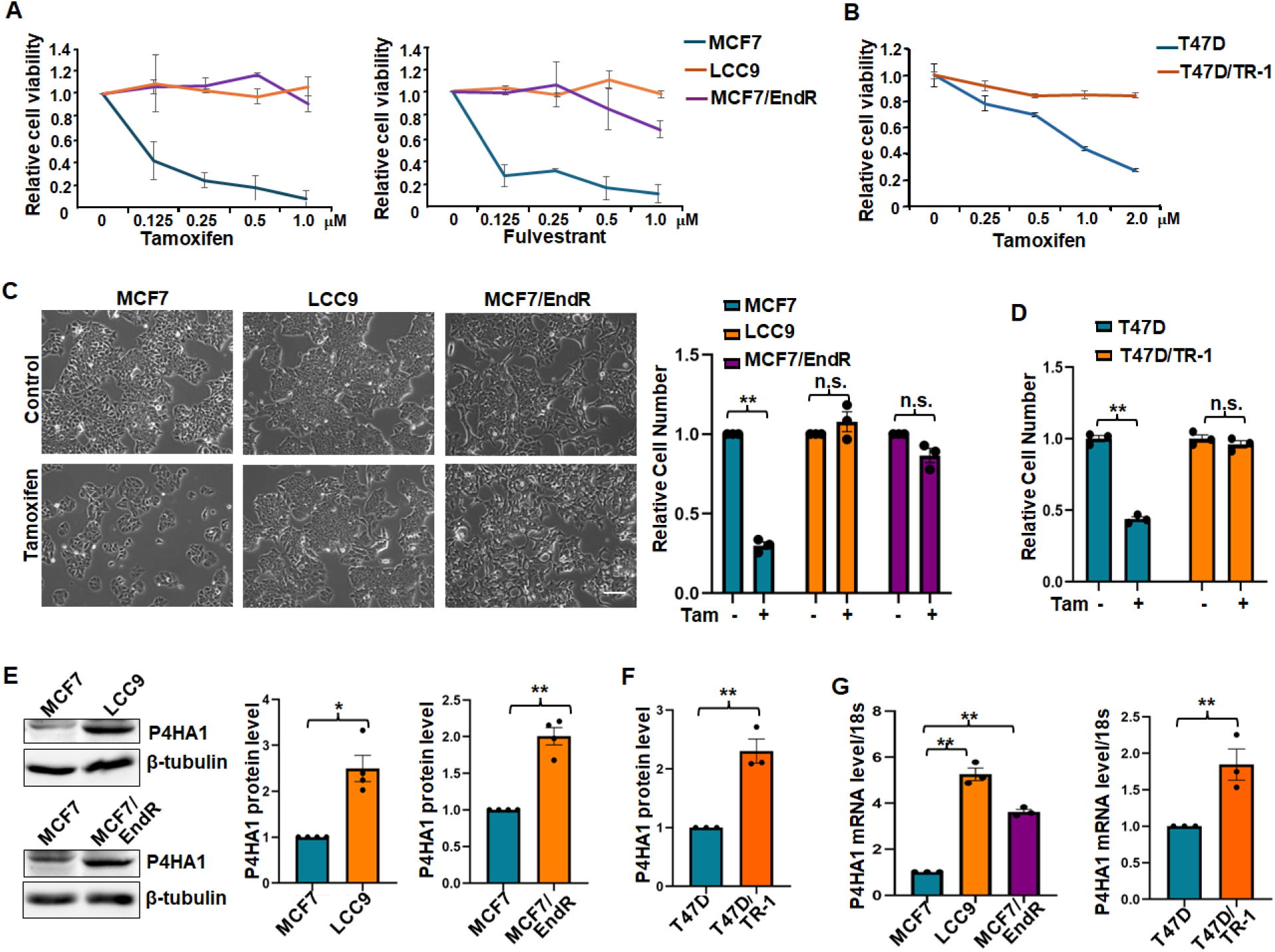
P4HA1 expression is associated with endocrine resistance. (A) CCK8 assay detected cell viability of parental MCF7cells, MCF7 variants LCC9 cells, and MCF7/EndR cells treated with tamoxifen or fulvestrant for 4 days. n=4. (B) Quantification of cell proliferation in parental T47D cells, T47D variant T47D/TR-1 cells treated with tamoxifen for 4 days. n=3. (C) Representative phase images and quantification of MCF7, LCC9 and MCF7/EndR cells treated with/without 1 μM tamoxifen. Scare bar: 50 μm. n=3. (D) Quantification of T47D and T47D/TR-1 cells treated with/without 1 μM tamoxifen. n=3. (E) Western blot analysis for expression of P4HA1 protein in parental MCF7 cells, endocrine-resistant LCC9 cells and MCF7/EndR cells. n=4. (F) Quantification expression of P4HA1 protein in parental T47D cells and tamoxifen-resistant T47D/TR-1 cells. n=3. (G) RT-qPCR analysis for expression of P4HA1 mRNA in parental MCF7 cells, endocrine-resistant LCC9 cells and MCF7/EndR cells; parental T47D cells, tamoxifen-resistant T47D/TR-1 cells. n=3. Results are presented as mean ± SEM, * p < 0.05, ** p < 0.01, n.s., no significant difference, independent Student’s t test for comparisons between two groups, one-way ANOVA test for comparisons among multiple groups. Tam: tamoxifen.

### Targeting P4HA1 restores sensitivity to endocrine therapy *in vitro*

To investigate the role of P4HA1 in modulating hormone therapy sensitivity *in vitro*, we established stable P4HA1 knockdown in endocrine-resistant luminal breast cancer cells. LCC9, MCF7/EndR, and T47D/TR-1 cells were infected with vector control or two distinct sh*P4HA1* lentiviral vectors encoding short hairpin RNA targeting *P4HA1* (sh*P4HA1*-1. sh*P4HA1*-2), both of which successfully silenced *P4HA1* expression (Fig. 3*A,* 3*D*, Fig. S3*A*). Upon treating the vector control or two *P4HA1*-silenced luminal breast cancer cells with endocrine agents, we observed that depleting P4HA1 restored tamoxifen sensitivity in resistant cell lines relative to controls (Fig. 3*B,* 3*C,* 3*E,* 3*F*). Because knockdown of P4HA1 via sh*P4HA1*-2 lentivirus alone resulted in slower baseline cell growth, sh*P4HA1*-1-silenced cells were selected for subsequent experiments. Next, we evaluated the sensitivity of *P4HA1*-silenced LCC9, MCF7/EndR cells to fulvestrant. The results showed that silencing *P4HA1* led to a significant reduction in cell growth following fulvestrant treatment (Fig. 3*G*, 3*H*, Fig. S3*B,* S3*C*). Furthermore, the *P4HA1*-silenced endocrine-resistant luminal breast cancer cells exhibited markedly reduced cell proliferation in response to endocrine treatment compared to control cells (Fig. 3*I-*3*L,* Fig. S3*D,* S3*E*). These data indicate that *P4HA1* silencing re-sensitizes endocrine-resistant luminal breast cancer cells to hormone therapy, demonstrating that P4HA1 expression is essential for maintaining the resistant phenotype. To test if P4HA1 expression is sufficient to drive resistance, we performed gain-of-function studies. Ectopic expression of P4HA1 failed to induce endocrine resistance in sensitive parental MCF7 cells (Fig. S4*A*- S4*C*), but its overexpression did promote partial tamoxifen resistance in sensitive parental T47D cells, without affecting fulvestrant sensitivity (Fig. S4*D*, S4*E*). These gain-of-function studies indicate that the ability of P4HA1 to drive endocrine resistance may depend on the specific genetic background of the host cell or the type of hormone therapy used. Taken together, these data indicate that while P4HA1 is functionally necessary to maintain resistance, its upregulation alone is insufficient to universally confer resistance in sensitive cells.

**Figure 3.**
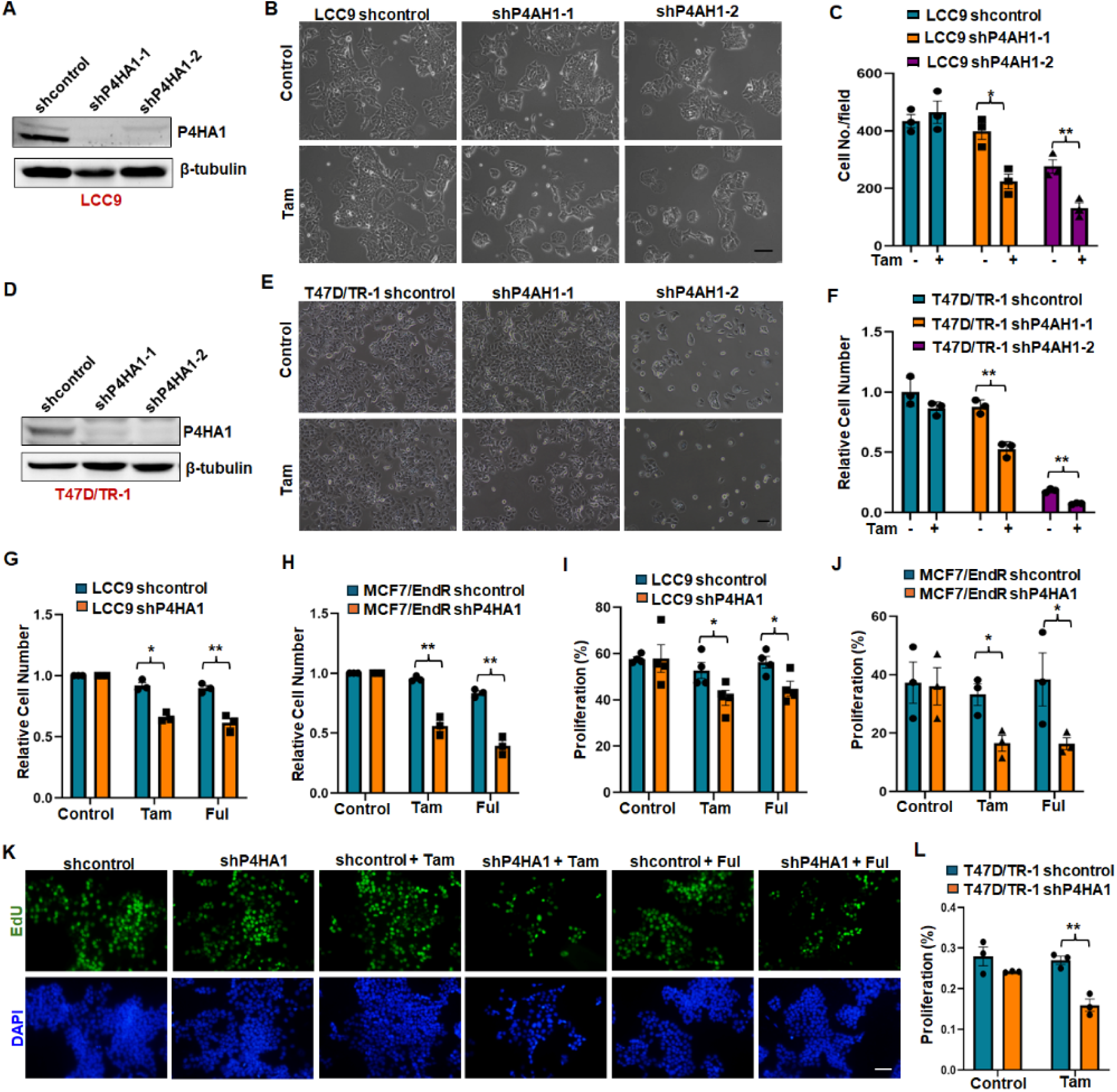
P4HA1 depletion restores sensitivity to endocrine treatment. (A) Western blot analysis knockdown efficiency of sh*P4HA1* lentivirus in LCC9 cells. (B) Phase images and (C) quantification of relative cell numbers of shcontrol and two sh*P4HA1* LCC9 cells with/without 1 μM tamoxifen treatment for 4 days. Scale bar: 50 μm. n=3. (D) Western blot analysis knockdown efficiency of sh*P4HA1* lentivirus in T47D/TR-1 cells. (E) Phase images and (F) quantification of relative cell numbers of shcontrol and sh*P4HA1* T47D/TR-1 cells with/without 1 μM tamoxifen treatment for 4 days. Scale bar: 50 μm. n=3 (G, H) Quantification of relative cell numbers in shcontrol and sh*P4HA1* LCC9 (G) and MCF7/EndR (H) cells with/without fulvestrant, tamoxifen for 4 days. n=3. (I) Quantification of cell proliferation in shcontrol and sh*P4HA1* LCC9 cells with/without fulvestrant, tamoxifen treatment for 4 days. n=4. (J) Quantification of cell proliferation in shcontrol and sh*P4HA1* MCF7/EndR cells with/without fulvestrant, tamoxifen treatment for 4 days. n=3. (K) EdU staining images of cell proliferation in shcontrol and sh*P4HA1* LCC9 cells with/without fulvestrant, tamoxifen treatment for 4 days. Scale bar: 25 μm. (L) Quantification of cell proliferation in shcontrol and sh*P4HA1* T47D/TR-1 cells with/without tamoxifen treatment for 4 days. n=3. Results are presented as mean ± SEM, *, P < 0.05; **, P < 0.01, independent Student’s t test. Tam: tamoxifen; Ful: fulvestrant.

We next evaluated the therapeutic potential of targeting *P4HA1* pharmacologically using the P4H inhibitors, ethyl-3, 4-dihydroxybenzoic acid (DHB) and 1, 4-dihydrophenonthrolin-4-one-3- Carboxylic acid (1, 4-DPCA) [23, 24]. Firstly, we determined the optimal, non-toxic treatment concentrations of DHB and 1, 4-DPCA in endocrine-resistant luminal breast cancer cell lines (Fig. S5*A*, S5*B*). Combining tamoxifen or fulvestrant with these P4H inhibitors significantly reduced cell growth in both LCC9 and MCF7/EndR (Fig. 4*A*-4*C,* Fig. S5*C,* S5*D*). Consistently, the combination of tamoxifen with either DHB or 1, 4-DPCA significantly decreased cell growth in T47D/TR-1 cells (Fig. 4*G*, Fig. S5*E*). This suppression was associated with a marked decrease in cell proliferation compared to treatment with any single agent alone (Fig. 3*D*-3*F*, 3*H*, Fig. S5*F*, S5*G*). Together, these results provide further evidence that P4HA1 is critical for maintaining endocrine resistance and demonstrate that its pharmacological inhibition partially restores sensitivity to endocrine therapy in resistant luminal breast cancer cells.

**Figure 4:**
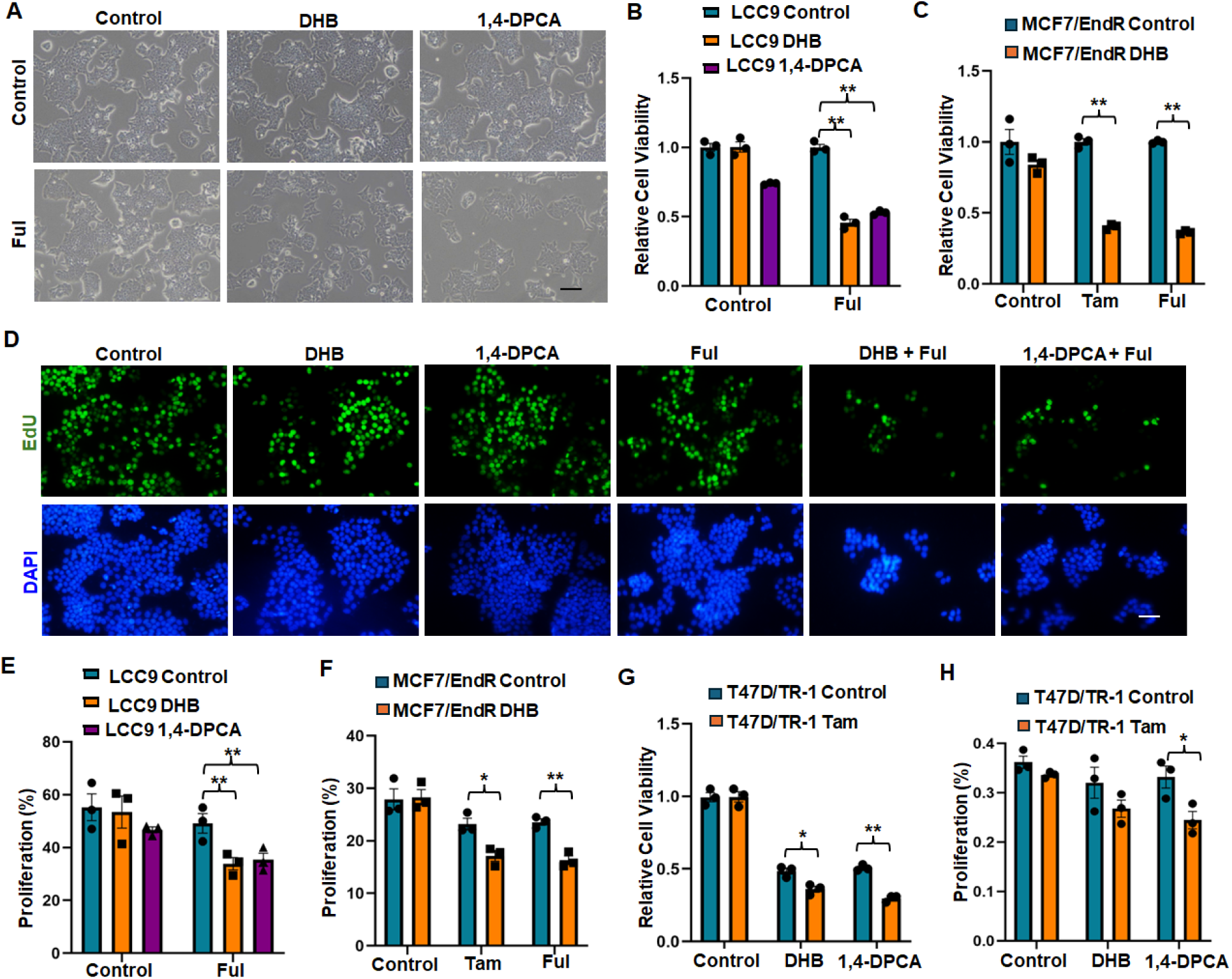
P4HA1 inhibition restores sensitivity to endocrine treatment. (A) Phase images of LCC9 cells treated with P4HA1 inhibitors (5 μM 1, 4-DPCA, 20 μM DHB) with/without 1 μM fulvestrant for 4 days. Scale bar: 50 μm. (B, C) Quantification of relative cell numbers in LCC9 (B), MCF7/EndR (C) cells treated with P4HA1 inhibitors (5 μM 1, 4-DPCA, 20 μM DHB), with/without 1 μM fulvestrant or 1 μM tamoxifen for 4 days. n=3. (D, E) EdU staining images and quantification of cell proliferation in LCC9 cells treated P4HA1 inhibitors (1, 4-DPCA, DHB) with/without 1 μM fulvestrant for 4 days. Scale bar: 25 μm. n=3. (F) Quantification of cell proliferation in MCF7/EndR cells treated P4HA1 inhibitor with/without 1 μM Tamoxifen, or fulvestrant for 4 days. n=3. (G) Quantification of relative cell numbers in T47D/TR-1 cells treated with P4HA1 inhibitors (5 μM 1, 4-DPCA, 15 μM DHB), with/without 1 μM tamoxifen for 4 days. n=3. (H) Quantification of cell proliferation in T47D/TR-1 cells treated P4HA1 inhibitors (1, 4-DPCA, DHB) with/without 1 μM Tamoxifen for 4 days. n=3. Results are presented as mean ± SEM, *, P < 0.05; **, P < 0.01, independent Student’s t test for comparisons between two groups, one-way ANOVA test for comparisons among multiple groups.

### Inactivation of P4HA1 re-sensitizes tumors to endocrine therapy in xenograft models

Our *in vitro* studies demonstrated that genetic silencing or pharmacological inhibition of P4HA1 effectively restored sensitivity to tamoxifen and fulvestrant in endocrine-resistant luminal breast cancer cells (Fig. 3, Fig. 4). To validate the role of P4HA1 in endocrine resistance *in vivo*, we evaluated whether P4HA1 inactivation could restore sensitivity to hormone therapy using xenograft mouse models. We orthotopically transplanted vector control or *P4HA1*-silenced LCC9 cells into the mammary fat pads of female SCID mice. While *P4HA1* knockdown alone did not affect tumor growth (Fig. 5*A,* 5*B*, Fig. S6*A*), fulvestrant administration significantly suppressed tumor progression in mice bearing *P4HA1*-silenced tumors compared to those with control tumors (Fig. 5*A*, *5B*, Fig. S6*A*). Consistent with *in vitro* data, immunohistochemical analysis of Ki67 in the harvested tumor tissues confirmed that this reduction in tumor volume was accompanied by a significant decrease in cellular proliferation (Fig. 5*C*, 5*D*).

**Figure 5:**
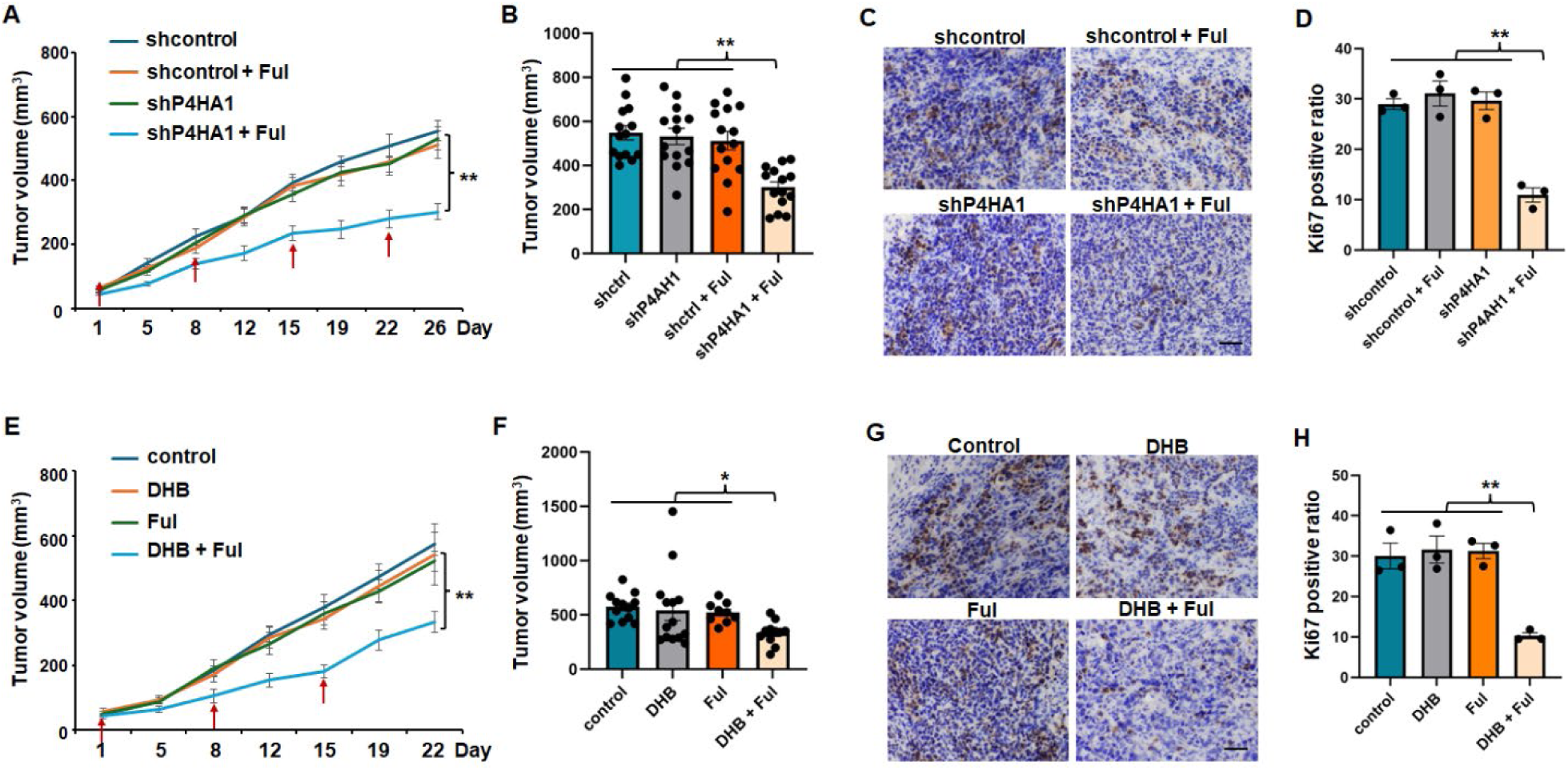
P4HA1 inactivation re-sensitizes endocrine-resistant luminal cancer to endocrine therapy *in vivo*. (A, B) Tumor growth curve and tumor volumes of control and *P4HA1*-silenced LCC9 cells implanted mice treated with control or fulvestrant. Red arrows mark fulvestrant administration. n=14. (C, D) IHC images of Ki67 staining and quantification of Ki67positive ratio in shcontrol and sh*P4HA1* LCC9 tumors collected from mice treated with control or fulvestrant. (E, F) Tumor growth curve and tumor volumes of LCC9 cells implanted mice treated with control, DHB, fulvestrant, DHB combination with fulvestrant. Red arrows mark fulvestrant administration. n=12 in control group, n=14 in DHB group, n=9 in fulvestrant group, n=11 in DHB plus fulvestrant group. (G, H) IHC images of Ki67 staining and quantification of Ki67positive ratio in LCC9 tumors collected mice treated with control, DHB, fulvestrant, DHB combination with fulvestrant. Scale bar: 50 μm. Results are presented as mean ± SEM, n=3, * p < 0.05, ** p < 0.01, one-way ANOVA test.

Next, we evaluated the therapeutic efficacy of combining fulvestrant with DHB, a P4HA1 inhibitor previously demonstrated to be safe in mouse [8]. In mice bearing parental LCC9 xenografts, the combination of DHB and fulvestrant significantly inhibited tumor growth and reduced final tumor weights compared to either agent alone (Fig.5*E*, 5*F*, Fig. S6*B*). Furthermore, this combination therapy significantly reduced the expression of cell proliferation marker Ki67 within the tumor tissues (Fig. 5*G*, 5*H*). Taking together, these findings demonstrate that P4HA1 inactivation, either through genetic silencing or pharmacological inhibition, effectively re-sensitizes endocrine-resistant tumors to hormone therapy *in vivo*. These results underscore the therapeutic potential of targeting P4HA1 as combination treatment strategies to overcome endocrine resistance in luminal breast cancer patients.

### P4HA1 modulates endocrine sensitivity via the ER stress pathway

Our *in vitro* and *in vivo* findings consistently indicate that targeting P4HA1 restores sensitivity to endocrine agents. To further elucidate the underlying molecular mechanisms, we performed RNA-seq analysis to compare gene expression patterns between control and *P4HA1*-silenced luminal breast cancer cells with endocrine treatment. DAVID functional annotation analysis revealed a significant enrichment of the endoplasmic reticulum (ER) stress signaling pathway in tamoxifen-treated P4HA1 knockdown cells (Fig. 6*A*, Fig. S7*A*). Specifically, the robust upregulation of ER stress-related genes suggests that P4HA1 depletion overcomes endocrine resistance by triggering an ER stress response. To validate these transcriptomic findings, we performed RT-qPCR analysis. The results confirmed that while *P4HA1* silencing or tamoxifen treatment alone modestly increased ER stress-related genes expression in resistant LCC9 cells, combining *P4HA1* knockdown with tamoxifen treatment induced the highest expression levels of ER stress-related genes (Fig. 6*B*).

**Figure 6:**
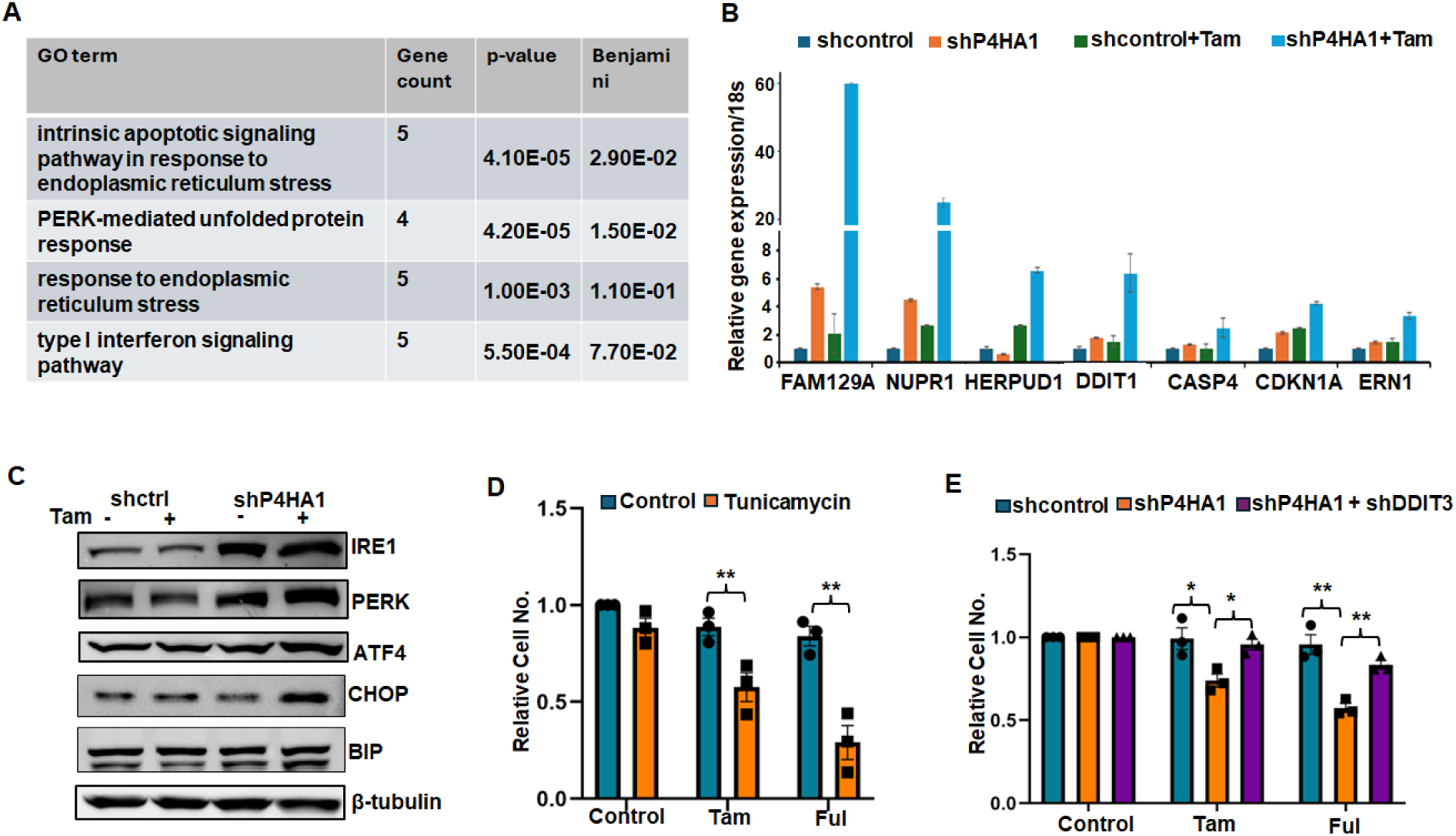
P4HA1 silencing combined with endocrine treatment activates ER stress signaling pathway. (A) DAVID analysis to identify the most significantly upregulated pathway in LCC9 sh*P4HA1* plus tamoxifen treatment group. (B) RT-qPCR analysis of differential gene expressions in shcontrol and sh*P4HA1* LCC9 cells with/without 1 μM tamoxifen treatment for 4 days. (C) Western blotting analysis of ER stress related molecules expression in shcontrol and sh*P4HA1* LCC9 cells with/without 1 μM tamoxifen treatment for 4 days. (D) Quantification of relative cell numbers in LCC9 treated with fulvestrant or tamoxifen treatment with/without 0.25 μg/mL tunicamycin. (E) Quantification of relative cell numbers in shcontrol LCC9 cells, sh*P4HA1* LCC9 cells with/without *DDIT3* silencing, treated with fulvestrant or tamoxifen treatment. Results are presented as mean ± SEM, n=3, * p < 0.05, ** p < 0.01, independent Student’s t test for comparisons between two groups, one-way ANOVA test for comparisons among multiple groups.

Protein Kinase R (PKR)-like ER Kinase (PERK) and Inositol-Requiring Enzyme 1 (IRE1) are key receptors in the ER stress-mediated signaling pathway [25, 26]. Activating Transcription Factor 4 (ATF4), binding immunoglobulin protein (BiP) (78 kDa glucose-regulated protein, GRP-78), and C/EBP homologous protein (CHOP) (DNA damage-inducible transcript 3, DDIT3) are key downstream mediators of this pathway [26] [27]. Therefore, we further evaluated the protein expression levels of these critical regulatory molecules. Western blotting results showed that the expressions of PERK, ATF4, and CHOP were elevated in *P4HA1*-silenced cells upon tamoxifen treatment (Fig. 6*C*). To determine whether inducing ER stress mimics the phenotype of P4HA1 inhibition, we treated endocrine-resistant luminal cancer with the canonical ER stress inducer tunicamycin [28, 29]. After optimizing the concentration of tunicamycin in LCC9 cells (Fig. S7*B,* S7*C*), we observed that pharmacological induction of ER stress with tunicamycin significantly re-sensitized resistant cells to endocrine treatment (Fig. 6*D*).

Furthermore, to confirm whether CHOP-mediated signaling functionally contributes to the cellular response, we performed rescue experiments by silencing *DDIT3* (Fig. S7*D,* S7*E*). Knockdown of *DDIT3* partially rescued cell growth, thereby reversing the endocrine-sensitizing phenotype observed in *P4HA1*-silenced luminal breast cancer cells (Fig 6E). CHOP /DDIT3 is a critical downstream pro-apoptosis transcription factor in the ER stress signaling pathway. Interestingly, Annexin V/PI flow cytometry evaluation revealed that neither *P4HA1* silencing nor treatment with P4H inhibitors did not trigger immediate physical execution of apoptosis at the evaluated time point (Fig. S7*F*, S7*G*). This suggests that P4HA1 inactivation induces an ER stress response that engages pro-apoptotic signaling pathways at the molecular level. This molecular activation can initially result in reduced proliferation prior to the onset of overt cell death, as observed in our study (Fig. 3, Fig. 4, and Fig. S7*F*, S7*G*.). Together, these data indicate that the loss of P4HA1 combination with endocrine therapy slows luminal breast cancer cell proliferation by activating an ER stress-mediated response.

### Inactivation of P4HA1 increases ROS levels during endocrine treatment

ER stress can be triggered by various physiological and pathological conditions [30–32], including the elevated levels of intracellular ROS [33–38]. ROS plays a role as double-edged sword in cancer. Moderately elevated ROS levels in tumor cells promote tumor progression. However, cancer cells with already elevated ROS and activated antioxidant systems are more sensitive to further increases in ROS [39]. Our previous work demonstrated that P4HA1 knockdown increases intracellular ROS levels and modulates oxidative phosphorylation in TNBC cells [8], therefore, we investigated whether P4HA1 similarly regulated ROS production in endocrine-resistant luminal breast cancer cells. We observed a significant increase of ROS accumulation in *P4HA1*- silenced endocrine-resistant luminal breast cancer cells under tamoxifen or fulvestrant treatment, compared to control cells (Fig. 7*A-*7*D*). Consistently, pharmacological inhibition of P4HA1 also significantly enhanced ROS levels when combined with endocrine agents compared to single-agent treatment groups (Fig. 7*E,* 7*F*, Fig. S8*A,* S8*B*).

**Figure 7:**
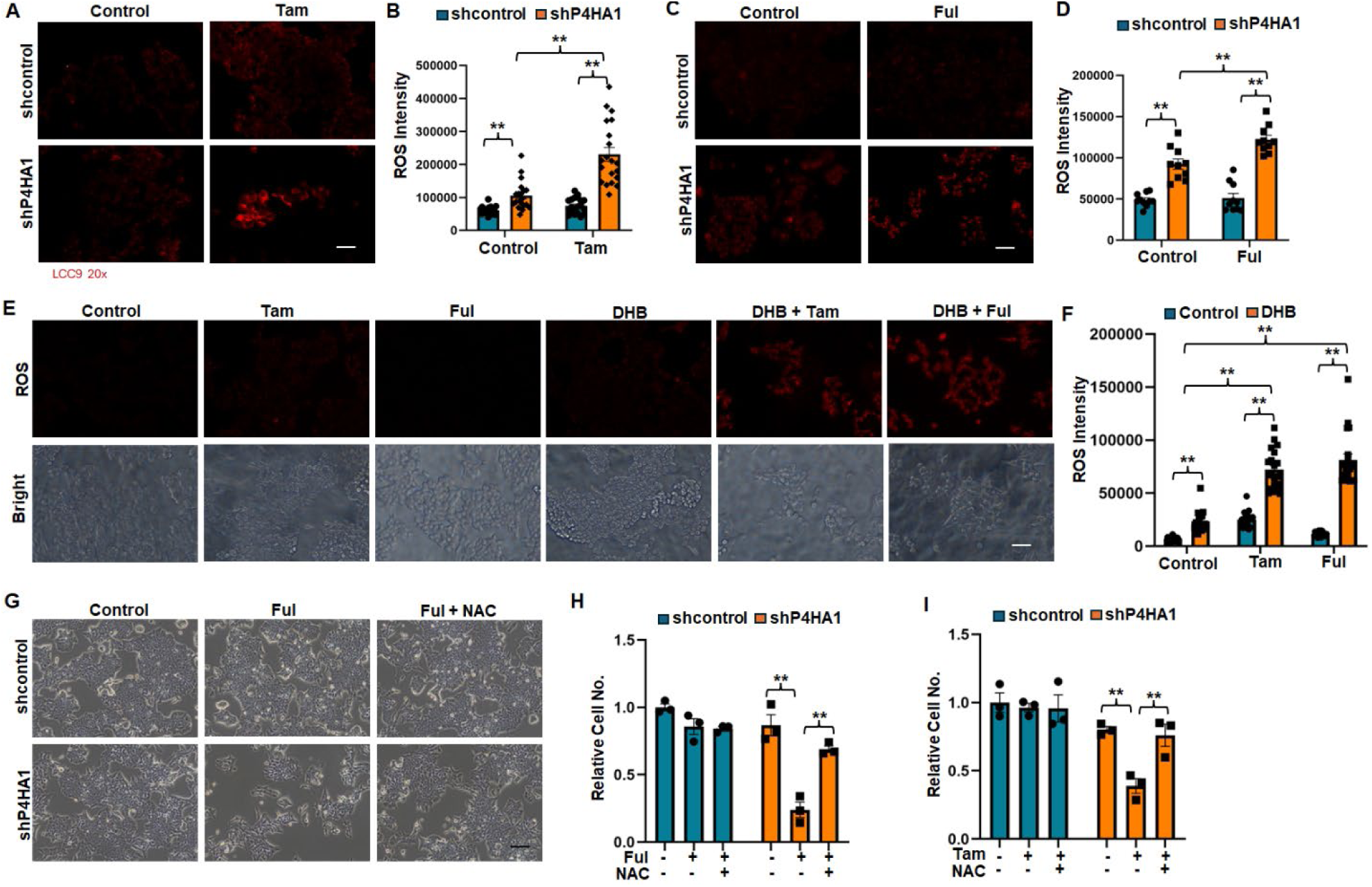
P4HA1 inactivation elevates cellular ROS levels upon endocrine treatment. (A, B) ROS staining images and quantification of ROS level in shcontrol and sh*P4HA1* LCC9 cells with/without 1 μM tamoxifen treatment for 4 days. (C, D) ROS staining images and quantification of ROS level in MCF7/EndR cells with/without 1 μM fulvestrant treatment for 4 days. (E, F) ROS staining images and quantification of ROS level in LCC9 tumors collected mice treated with control, DHB, fulvestrant, DHB combination with fulvestrant. (G) Phase images of shcontrol and sh*P4HA1* LCC9 cells with 1 μM fulvestrant with/without 1 μM NAC treatment for 4 days. (H, I) Quantification of relative cell number of shcontrol and sh*P4HA1* LCC9 cells with 1 μM fulvestrant or tamoxifen with/without 1 μM NAC treatment for 4 days. Scale bar: 25 μm. Results are presented as mean ± SEM, n=3, **, P < 0.01, one-way ANOVA test.

To further assess the functional relevance of ROS accumulation in P4HA1-modulating endocrine sensitivity, we treated P4HA1-knockdown cells with N-acetyl-L-cysteine (NAC), a commonly used ROS scavenger [40, 41]. Antioxidant treatment with NAC partially rescued cell growth, reversing the endocrine-sensitizing effects of *P4HA1*-silenced luminal breast cancer cells, indicating that elevated ROS levels play a crucial functional role in resensitizing resistant cells to endocrine therapy (Fig. 7*G-*7*I,* Fig. S8*C*). Together, these findings suggest that P4HA1 contributes to endocrine resistance in luminal breast cancer by suppressing oxidative stress, establishing a therapeutic vulnerability that can be exploited via P4HA1 inhibition.

## Discussion

Endocrine therapy has been utilized for decades and remains the first-line treatment for patients with estrogen receptor positive luminal breast cancer. Although it has significantly reduced mortality [10], the development of resistance to endocrine therapy and subsequent lethal recurrence remain primary clinical challenges. Elucidating the mechanisms driving both acquired and intrinsic (*de novo*) resistance is critical to improve outcomes of breast cancer patients. Over the past decades, researchers have uncovered various crucial mechanisms of endocrine resistance, including estrogen receptor downregulation, *ESR1* mutations or fusions, aberrant activation of growth signaling pathways, altered cell cycle progression, and diverse and complex changes in tumor microenvironment [42–44]. However, the mechanisms underlying endocrine resistance are complicated and remain partially undefined. Here, we report that a hydroxylation enzyme P4HA1 is highly expressed in endocrine-resistant luminal breast cancer cells and human luminal B breast tumors, where its elevated expression significantly correlates with shorter relapse-free survival in patients with luminal B breast cancer. Genetic silence or pharmacological inhibition of P4HA1 restored endocrine sensitivity in resistant luminal breast cancer cells, demonstrating that P4HA1 expression is essential for maintaining the resistant phenotype. A notable finding of this study is that P4HA1 ectopic expression is insufficient to universally confer resistance in sensitive parental cells; its overexpression induced partial tamoxifen resistance in parental T47D cells but failed to alter the sensitivity of MCF7 cells or affect responsiveness to fulvestrant. This context-dependent outcome suggests that P4HA1 likely required specific co-factors, downstream effectors, or a permissive genetic background to confer endocrine sensitivity. Our loss-of- and gain-of-function studies collectively demonstrate that P4HA1 is necessary for the maintenance of advanced endocrine resistance, but its ability to induce resistance in sensitive parental cells is highly constrained by the host cell’s genetic background and the specific endocrine therapy employed. Therapeutically, targeting P4HA1 remains a highly viable strategy to overcome established endocrine resistance.

The endoplasmic reticulum (ER) is a critical cellular organelle responsible for protein synthesis, folding, modification, and transportation [45]. The canonical role of P4HA1 is to hydroxylate collagen and other substrates within the ER, a modification essential for proper protein folding and secretion. Accumulation of unfolded or misfolded proteins in the ER lumen initiates the unfolded protein response (UPR), a stress response aimed at restoring cellular homeostasis. Our findings demonstrate that *P4HA1* silencing triggers ER stress in endocrine-resistant breast cancer cells; combing P4HA1 depletion with endocrine therapy exacerbates ER stress to cytotoxic levels, thereby suppressing cell proliferation and restoring sensitivity to endocrine therapy. While our acute experimental timeline captures significant activation of pro-apoptotic signaling pathways at molecular level, no immediate induction of physical apoptosis was observed. Therefore, we cannot rule out an initial contribution of a transient growth arrest prior to cell death. Future longer-term studies will be essential to delineate the definitive long-term fate of these cells. These observations align with recent studies reporting up-regulation of the ER stress pathway in endocrine-resistant, hormone receptor-positive breast cancer [46, 47]. Notably, Pal and colleagues demonstrated that endocrine-resistant breast cancer cells exhibit increased sensitivity to ceramide-induced lethal ER stress through PERK pathway activation [46]. Together, these findings highlight the ER stress pathway as a targetable vulnerability in endocrine-resistant breast cancer. Accumulation of intracellular ROS, when elevated beyond a critical threshold, can trigger overwhelming ER stress [33–38]. Conversely, persistent ER stress can accelerate ROS production, creating a feed-forward loop of cellular distress [36]. Sustained ER stress and oxidative stress drive the pathogenesis of various human disorders [48]. Thus, P4HA1-mediated regulation of ER stress and ROS accumulation may synergistically promote endocrine resistance. Our findings uncover a previously unrecognized role for P4HA1 in breast cancer progression, in which it coordinates the critical crosstalk between endocrine resistance and cellular stress pathways.

Canonically, P4HA1 hydroxylates proline into 4-hydroxyproline at the Y position of the collagen -X-Y-Gly- triplet motif, which is essential for the mature collagen trimer stability and secretion [1] [2]. However, we did not observe significantly increased collagen deposition in endocrine-resistant luminal breast cancer cells compared with endocrine-sensitive luminal breast cancer cells (unpublished data). This suggests that the highly expressed P4HA1 in endocrine-resistant luminal breast cancer operates through a collagen-independent manner. Our prior work demonstrated that P4HA1 promotes chemoresistance in TNBC by reducing proline hydroxylation on hypoxia-inducible factor (HIF) 1α, consequently modulating HIF1α-dependent cancer cell stemness [8]. Literature indicates that hyperactivation of HIF1α drives tamoxifen resistance and enhances aerobic glycolysis through modulation of Akt/mTOR and/or AMPK signaling pathways [49]. Compared to endocrine-sensitive breast cancer cells, endocrine-resistant cells contain a higher proportion of cancer stem cells [50], exhibit activated HER2-associated signaling molecules (pAKT and pERK), and express mesenchymal biomarkers [51]. As shown in this study and our previous work, P4HA1 is highly expressed in HER2-positve and TNBC breast tumors. These evidence from the literature suggest that in luminal breast tumor, P4HA1 may contribute to multifactorial mechanisms underlying endocrine resistance, potentially through regulation of HIF1α signaling, cancer stemness, glycolysis, and epithelial–mesenchymal transition (EMT). Fully elucidating these mechanisms remains an important direction for future investigation.

Although targeted inhibitors against PI3K, mTOR, and CDK4/6 have been approved by the FDA to improve outcomes for patients with endocrine-resistant breast cancer, a substantial proportion of patients still fail to respond or ultimately relapse. Given that the molecular mechanisms underlying endocrine resistance are manifold, developing optimal combination regimens is essential to delay cancer adaptation to treatment and prevent secondary resistance [43]. Our results demonstrate that either genetic silence of *P4HA1* using shRNA or pharmacological inhibition of P4HA1 with small molecule inhibitors effectively restores sensitivity to endocrine therapy in endocrine-resistant breast cancer cell models. As a high-priority direction of our study will focus on dual endocrine/CDK4/6 inhibitor resistance, we will establish independent fulvestrant/CDK4/6 inhibitor-resistant models for future study. Our findings highlight P4HA1 as a promising therapeutic target for advanced luminal breast cancer. Consequently, evaluating P4HA1 inhibition as a part of combination treatment strategies, particularly for patients with high P4HA1 expression, presents a compelling translational approach to overcome therapy resistance and improve long-term clinical outcomes.

## Abbreviations

P4HA1: alpha subunit of Prolyl 4-hydroxylase;
SERMs: selective estrogen receptor modulators;
SERDs: selective estrogen receptor degraders;
PROTACs: proteolysis target chimeras;
P4H: Prolyl 4-hydroxylase;
P4HB: beta subunit of Prolyl 4-hydroxylase;
ER: endoplasmic reticulum;
ROS: reactive oxygen species;
UPR: unfolded protein response;
DMEM: Dulbecco’s Modified Eagle’s Medium;
FBS: fetal bovine serum;
TMA: tumor microarray;
DHB: ethyl-3, 4-dihydroxybenzoic acid;
1,4-DPCA: 1, 4-dihydrophenonthrolin-4-one-3-Carboxylic acid;
PERK: Protein Kinase R (PKR)-like ER Kinase,
IRE1: Inositol-Requiring Enzyme 1; ATF4, Activating Transcription Factor 4;
BIP: Binding immunoglobulin protein;
GRP-78: 78 kDa glucose-regulated protein
CHOP: C/EBP homologous protein;
DDIT3: DNA damage-inducible transcript 3;
NAC: N-acetyl-L-cysteine;
HIF1α: hypoxia-inducible factor 1α
EMT: epithelial–mesenchymal transition
PDI: protein disulfide isomerase.

## Declarations

### Availability of data and materials

All data generated or analyzed during this study are included in this published article and its supplementary information files.

### Competing interests

The authors declare that they have no competing interests.

### Funding

This work is supported by American Cancer Society (IRG-16-182-28), National Institutes of Health (R21AG080176), the start-up funding from OSU Comprehensive Cancer Center to G.X, and National Institutes of Health (R01CA207772, R01CA277946, and R01CA274981) to R.X.

### Authors’ contributions

G.X. conceived the work, performed, and supervised the data analysis, and wrote the manuscript.

D.H. and B.K. performed the experiments, collected, and analyzed the data, and edited the manuscript. R.X. analyzed the data and edited the manuscript. D.K. and H.L. edited the manuscript. R.S., provided reagents and edited the manuscript. All authors have approved the manuscript for publication.

## Acknowledgements

The authors acknowledge the assistance of Markey Cancer Center Shared Resource Facilities, all of which are supported by the grant P30 CA177558: the Biospecimen and Tissue Procurement Shared Resource Facility for assistance in tissue fixation; the Flow Cytometry and Cell Sorting Core Facility for performing FACS analysis. DK is supported by the NIH/NCI 1K22CA292325-01A1, Lung Cancer Research Foundation, American Lung Association-INALA2026, and OSU start-up. The human breast cancer cell line LCC9 is a gift from Dr. Robert Clarke, Georgetown University.

## Notes

### Competing Interest Statement

The authors have declared no competing interest.

