## Supplemental figures for "P4HA1 sustains endocrine resistance in luminal breast cancer via regulation of cellular stress"

**Supplemental Figure 1:**

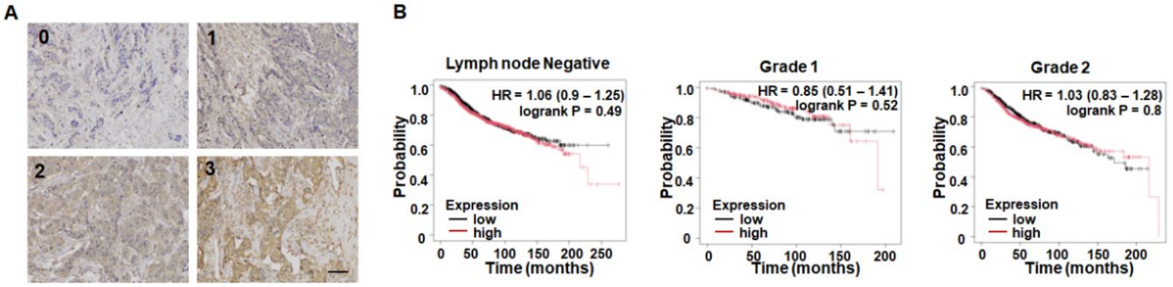


**Figure S1**: P4AH1 staining images from breast cancer TMA and Kaplan Meier analysis in breast cancer patients. (A) P4HA1 IHC staining scores 0-3; (B) Kaplan Meier analysis of survival in lymph node negative (n=2277) and grade 1 (n=397), grade 2 (n=1177) breast cancer patients with different levels of P4HA1 protein levels.

**Supplemental Figure 2:**

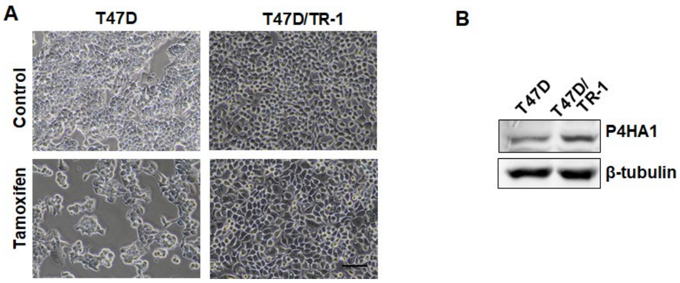


**Figure S2**: Upregulated expression of P4HA1 in tamoxifen-resistant cells. (A) Phase images of parental T47D and tamoxifen-resistant T47D/TR-1 cells with/without 1 μM tamoxifen treatment for 4 days. 50 μm. (B) Western blot analysis for expression of P4HA1 protein in parental T47D cells and tamoxifen-resistant T47D/TR-1 cells.

**Supplemental Figure 3:**

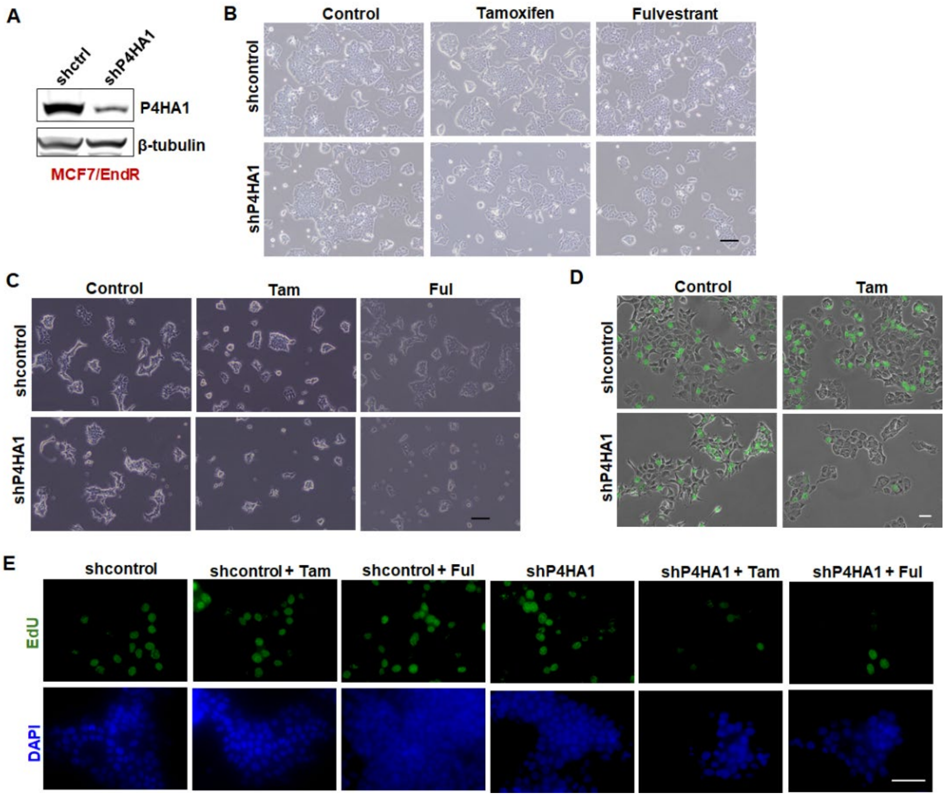


**Figure S3**: Silencing P4AH1 restores endocrine sensitivity *in vitro*. (A) Western blot analysis knockdown efficiency of sh*P4HA1* lentivirus in MCF7/EndR cells. (B) Phase images and quantification of shcontrol and sh*P4HA1* LCC9 cells with/without 1 µM tamoxifen or fulvestrant treatment for 4 days. n=3. Scare bar: 50 µm. (C) Phase images of shcontrol and sh*P4HA1* MCF7/EndR cells with/without 1 µM tamoxifen or fulvestrant treatment for 4 days. Scare bar: 50 µm. (D) Representative EdU staining images of shcontrol and sh*P4HA1* T47D/TR-1 cells treated with/without 1 µM tamoxifen treatment for 4 days. Scare bar: 25 µm. (E) Representative EdU staining images of shcontrol and sh*P4HA1* MCF7/EndR cells treated with/without 1 µM tamoxifen or fulvestrant treatment for 4 days. Scare bar: 25 µm.

**Supplemental Figure 4:**

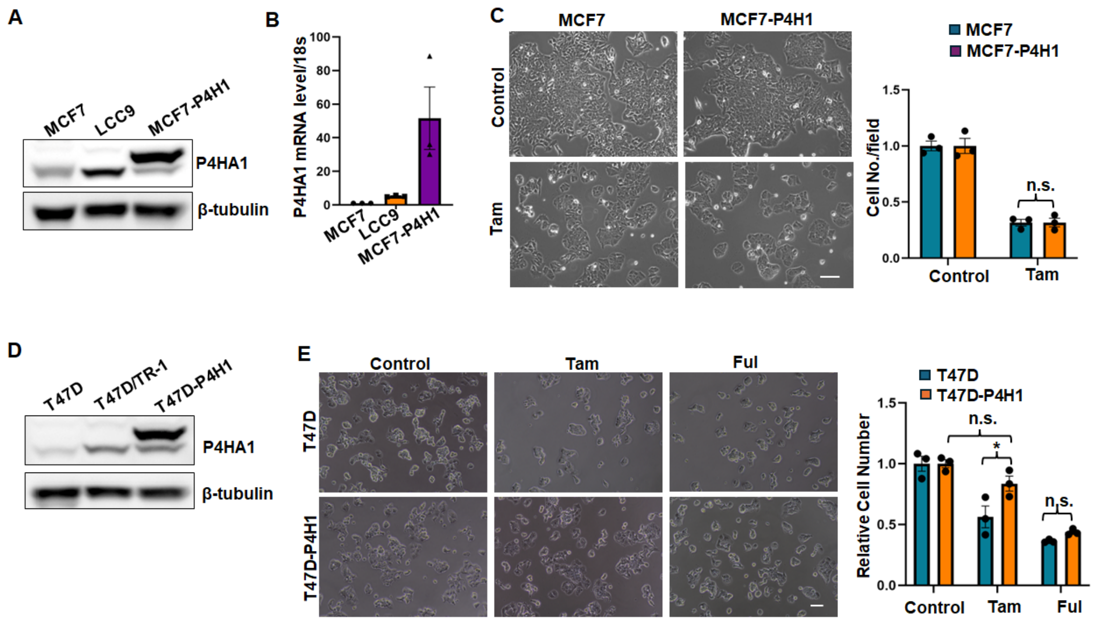


**Figure S4**: Exogenous overexpression of P4HA1 in sensitive parental cells. (A, B) Western blot and RT-qPCR confirmation of P4HA1 overexpression in MCF7-P4H1 cells. (C) Phase images and quantification of relative cell numbers in MCF7 and MCF7-P4H1 cells with/without 1 μM tamoxifen treatment. (D) Western blot confirmation of P4HA1 overexpression in T47D-P4H1 cells. (E) Phase images and quantification of relative cell numbers in T47D and T47D-P4H1 cells with/without 1 μM tamoxifen or fulvestrant treatment. Scale bar: 50 μm. Results are presented as mean ± SEM. n=3. *, $P < 0.05$, **, $P < 0.01$, n.s., no significant difference, independent Student's t test for comparisons between two groups, one-way ANOVA test for comparisons among multiple groups.

**Supplemental Figure 5:**

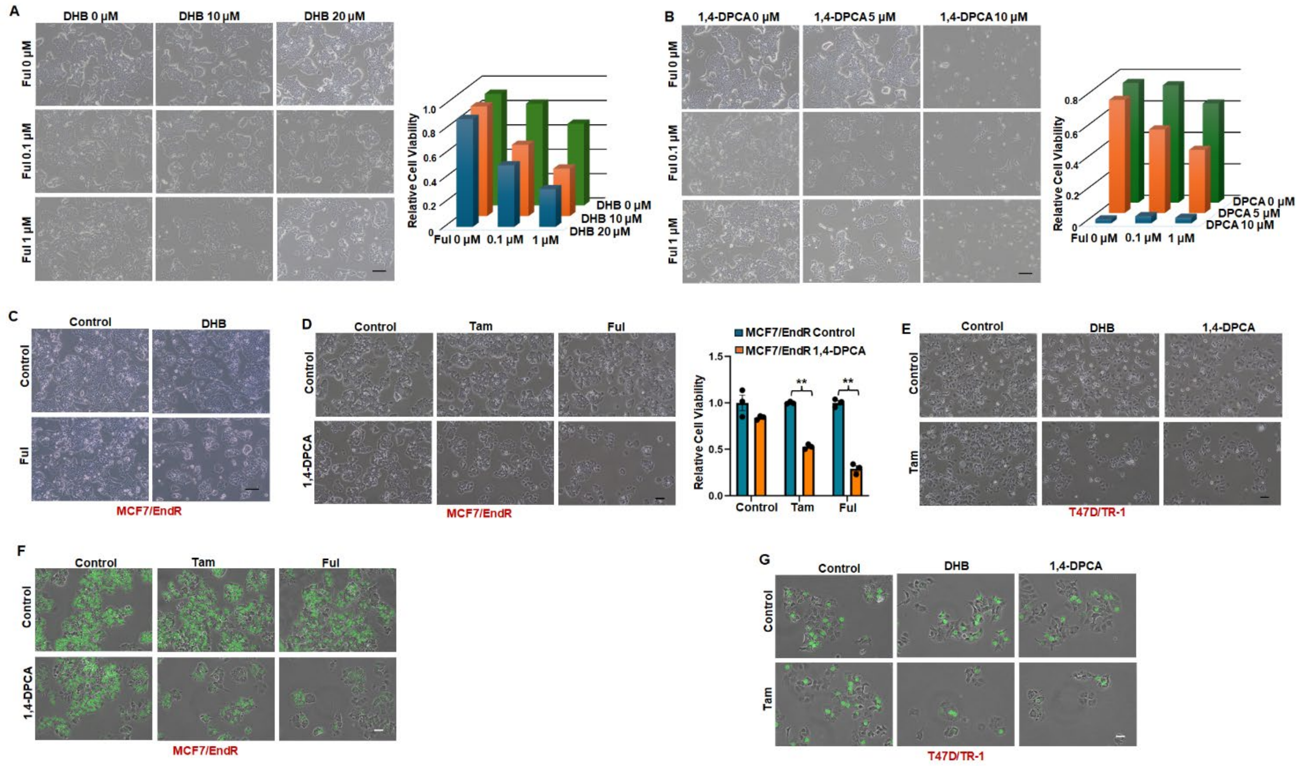


**Figure S5**: Pharmacological inhibition of P4AH1 restores endocrine sensitivity *in vitro*. (A) Representative phase images and quantification of LCC9 cells treated different concentrations of DHB (0, 10 μM, 20 μM) and fulvestrant (0, 0.1 μM, 1 μM). Scare bar: 50 μm. (B) Representative phase images and quantification of LCC9 cells treated different concentrations of 1,4-DPCA (0, 5 μM, 10 μM) and fulvestrant (0, 0.1 μM, 1 μM). Scare bar: 50 μm. (C) Representative phase images of MCF7/EndR cells treated with 20 μM DHB with/without 1 μM fulvestrant. Scare bar: 50 μm. (D) Representative phase images and quantification of MCF7/EndR cells treated with 5 μM 1,4-DPCA with/without 1 μM tamoxifen or 1 μM fulvestrant. Scare bar: 50 μm. Results are presented as mean ± SEM. **, $P < 0.01$, independent Student's t test. (E) Representative phase images of T47D/TR-1 cells treated with 15 μM DHB or 5 μM 1,4-DPCA with/without 1 μM tamoxifen. Scare bar: 50 μm. (F) Representative EdU staining images of MCF7/EndR cells treated with 5 μM 1,4-DPCA with/without 1 μM tamoxifen or 1 μM fulvestrant. Scare bar: 25 μm. (G) Representative EdU staining images of T47D/TR-1 cells treated with 15 μM DHB, or 5 μM 1,4-DPCA with/without 1 μM tamoxifen. Scare bar: 25 μm.

combined treatment for 4 days. (K) EdU staining images and quantification proliferation of LCC9/PalR cells treated with P4HA1 inhibitor (20 μM DHB) with/without fulvestrant and palbociclib combined treatment for 4 days. Scale bar: 50 μm. Results are presented as mean ± SEM. n=3. *, $P < 0.05$, **, $P < 0.01$, n.s., no significant difference, independent Student's t test for comparisons between two groups, one-way ANOVA test for comparisons among multiple groups.

**Supplemental Figure 6:**

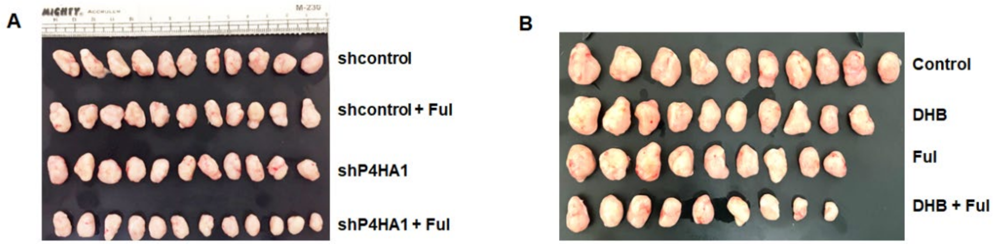


**Figure S6**: Inactivation of P4AH1 restores endocrine sensitivity *in vivo*. (A) Tumors of control and *P4HA1*-silenced LCC9 cells implanted mice treated with control or fulvestrant. (B) Tumors of LCC9 cells implanted mice treated with control, DHB, fulvestrant, DHB combination with fulvestrant.

**Supplemental Figure 7:**

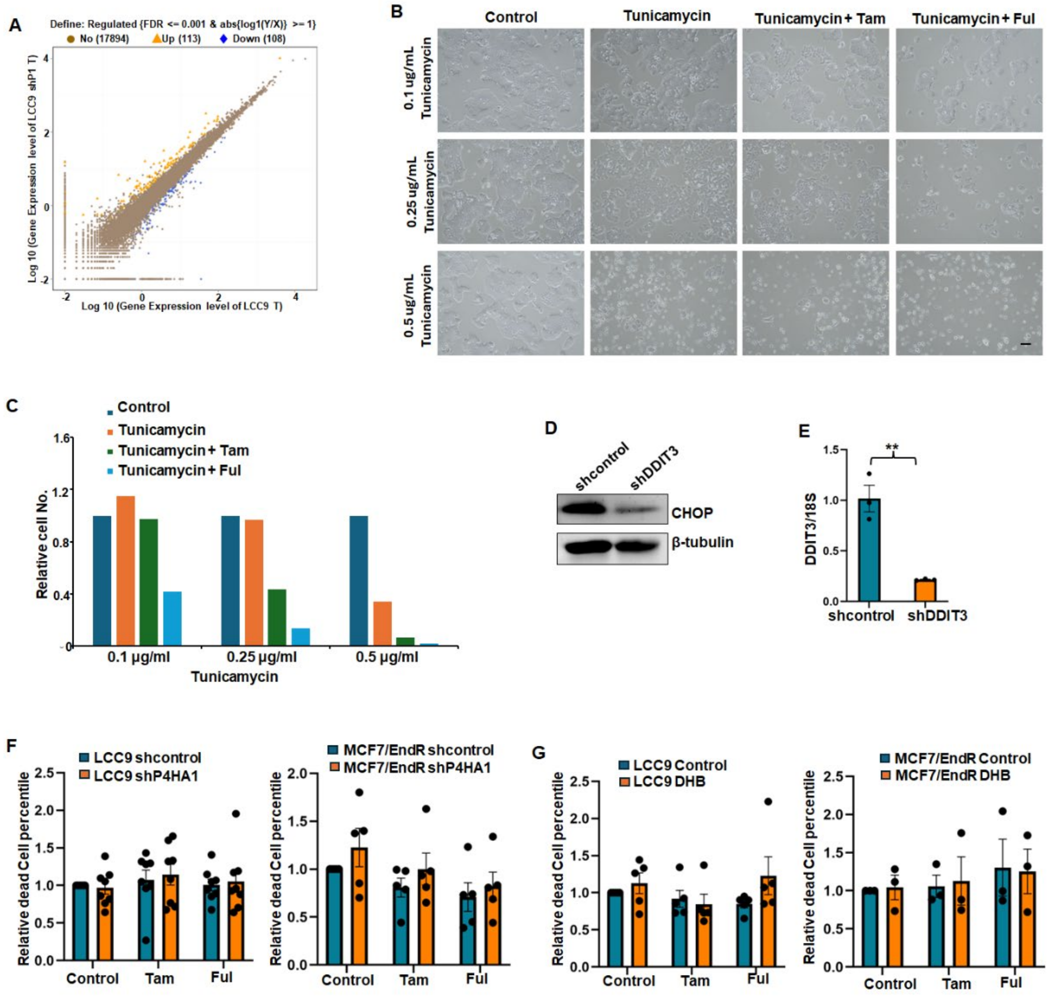


**Figure S7**: ER stress inducer treatment and CHOP silencing in LCC9 cells. (A) Differential gene expression analysis scatter plot comparing LCC9 T (vector control LCC9 treated with tamoxifen) and LCC9 shP1 T (sh*P4AH1* LCC9 treated with tamoxifen) samples. This figure presents a scatter plot where each point represents a gene, illustrating the correlation and differential expression levels between two conditions, using a log10 expression scale. (B, C) Representative phase images and quantification of LCC9 cells treated different concentrations of tunicamycin (0.1 µg/mL, 0.25 µg/mL, 0.5 µg/mL) combination with 1 µM 4OHT or fulvestrant. (D, E) Western blot and RT-PCR analysis knockdown efficiency of sh*DDIT3* lentivirus in LCC9 cells. Results are presented as mean ± SEM, n=3, **, $P < 0.01$, independent Student's t test. (F) Quantification of cell apoptosis in shcontrol and sh*P4HA1* LCC9 cells and MCF7/EndR cells with/without fulvestrant, tamoxifen treatment for 4 days via Annexin V & PI staining analysis. Results are presented as mean ± SEM. (G) Quantification of cell apoptosis in LCC9 cells and MCF7/EndR cells treated with P4HA1 inhibitors DHB with/without fulvestrant, or tamoxifen via annexin V & PI staining analysis. Results are presented as mean ± SEM.

**Supplemental Figure 8:**

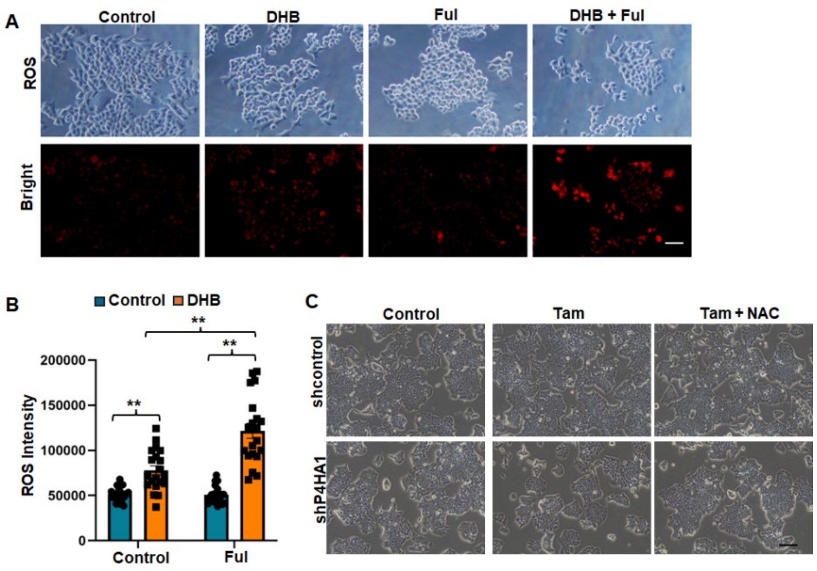


**Figure S8**: ROS staining and ROS inhibitor NAC treatment in LCC9 cells. (A, B) ROS staining images and quantification of ROS levels in MCF7/EndR cells treated with control, DHB, fulvestrant, DHB combination with fulvestrant. Scale bar: 25 µm. Results are presented as mean ± SEM, n=3, **, P < 0.01, independent Student's t test. (C) Phase images of shcontrol and sh*P4HA1* LCC9 cells with 1 µM 4OHT with/without 1 µM NAC treatment for 4 days. Scale bar: 50 µm.
